# Maternal Effect Senescence and Fitness: A Demographic Analysis of a Novel Model Organism

**DOI:** 10.1101/847640

**Authors:** Christina M. Hernández, Silke F. van Daalen, Hal Caswell, Michael G. Neubert, Kristin E. Gribble

## Abstract

Maternal effect senescence—a decline in offspring fitness with maternal age—has been demonstrated in a range of taxa, including humans. Despite decades of phenotypic studies, it remains unclear how maternal effect senescence impacts population structure or evolutionary fitness. To understand the impact of maternal effect senescence on population dynamics, fitness, and selection, we used data from individual-based culture experiments on the microscopic aquatic invertebrate, *Brachionus manjavacas* (Rotifera), to develop a series of matrix population models in which individuals are classified jointly by age and maternal age. By comparing the results derived from models with and without maternal effects, we found that the fitness difference due to maternal effect senescence arises primarily through decreased fertility, particularly at maternal ages corresponding to the peak reproductive output. In all models, selection gradients decrease with increasing age. They also decrease with maternal age for large maternal ages, implying that maternal effect senescence can evolve through the same process as in Hamilton’s theory of the evolution of demographic senescence. We find that maternal effect senescence significantly alters population structure and fitness for *B. manjavacas*, a species with high maternal investment and maximum reproduction in early-to mid-life. The models developed here were built with data from an emerging model organism, and are widely applicable to evaluate the fitness consequences of maternal effect senescence across species with diverse aging and fertility schedule phenotypes.

## 1 Introduction

In many species, survival and reproduction decrease with advancing age, a process known as *senescence*. The evolution of senescence was one of the earliest questions in life history theory [1, 2] and has been studied extensively in the laboratory, in theory, and in the field (e.g., [3–6]). The evolution of senescence is explained by the age patterns of selection gradients, a measure of the strength of selection, defined as the derivative of the population growth rate with respect to a given trait value. Hamilton [2] showed that age-specific selection gradients on mortality and fertility decrease with age. Thus, traits expressed early in life have a larger impact on fitness than those expressed later. As a result, selection will favour traits that lead to negative effects on mortality and fertility at older ages if there are even small beneficial effects in youth.

*Maternal effect senescence* refers to reduced success or quality of offspring with advancing age of the mother [7]. Advanced maternal age has known negative effects on offspring health, lifespan and fertility in humans and other species [8–16]. In many taxa, including *Drosophila*, rotifers, seed beetles, *Daphnia*, and birds, offspring from older mothers have shorter lives, lower reproductive success, or both [8, 11, 14, 16–19]. In humans, advanced maternal age is associated with shorter lifespan [20] and decreased health of offspring, including increased incidence and earlier onset of neurodegenerative diseases [9, 21]. In *Caenorhabditis elegans, Daphnia*, and rotifers, advanced maternal age leads to increases in offspring size, changes in development time, and increased variability in gene expression [16, 22–24].

Maternal effect senescence poses a challenge to evolutionary theory. Hamilton’s [2] results are based on a model in which individuals are distinguished only by differences in their age. The description of maternal effect senescence makes it clear that age does not completely characterize an individual. Instead, the age of the individual and the age of its mother at the time of its birth combine to determine survival and reproduction. Thus, the calculation of fitness and selection gradients relating to maternal effect senescence requires a multistate, age-by-stage demographic model, where the “stage” refers to maternal age. It is known that such multistate models can fundamentally alter the patterns of age-specific selection gradients and the evolution of senescence [25].

In this paper, we develop such a model, producing a novel framework that is generally applicable to maternal effect senescence and to other non-Mendelian maternal effects that influence offspring survival and fecundity, including maternal diet, maternal stress, and inducible defenses [22, 26–32]. We analyze the model to document selection gradients on survival and fertility as functions of age and maternal age, and to quantify the fitness impact of maternal age effects throughout the life cycle.

We apply the model to the monogonont rotifer, *Brachionus manjavacas*, an ideal system in which to investigate maternal effects on offspring fitness [16, 33, 34]. *B. manjavacas* is an ecologically-important microscopic, aquatic, invertebrate animal with approximately 1000 cells and with distinct nervous, digestive, muscle, and reproductive systems. *Brachionus* is simple to culture in the laboratory and has a lifespan of two weeks, making it a tractable model system for life history and other demographic studies. Embryos carried by females are brooded externally, permitting easy collection of eggs and creation of age-synchronized cohorts. *Brachionus* rotifers can be individually housed and monitored, permitting high-frequency collection of individual-level measurements of lifespan and fecundity with high replication. Because reproduction is asexual, individuals within a population are genetically identical. This eliminates variability due to genetic recombination or paternal effects. *B. manjavacas* females produce 25–30 offspring sequentially and continuously over a reproductive period of approximately 10 days. Maternal investment in offspring is high: eggs and neonates measure nearly 1/3 the size of the mother. *Brachionus* rotifers have direct development, with no larval stage or metamorphosis, and exhibit no post-hatching parental care. Thus, changes in maternal age are not confounded with changes in maternal care. Although *Brachionus* rotifers have been used in previous studies of maternal effects, the impacts of maternal age effects on population dynamics and evolutionary fitness have not been investigated.

We begin by describing the demographic model, then describe the experimental system and the estimation of the survival and fertility parameters. The model provides estimates of fitness, stable population structure, and selection gradients on survival and fertility. We use life table response experiment (LTRE) methods [e.g. 35, 36] to decompose the impact of maternal effect senescence into contributions from age and stage-specific survival and fertility. We compare the results between models with and without such senescence in both high growth rate (laboratory) and low growth rate (pseudo-natural) environments. These contrasts are possible only within the context of a model such as that we develop here. Our results reveal that maternal age has a significant impact on population growth rate and age structure across environments, and have implications for the evolution of aging and of maternal age effects.

## 2 The demographic model

Our demographic model is a special case of a general age-by-stage structured model thoroughly described by Caswell et al. [37]. If *n*_*i,j*_(*t*) is the number of individuals in maternal age class *i* and age class *j* at time *t*, then the composition of the population is given by a column vector **ñ**(*t*), that collects maternal ages within age classes:

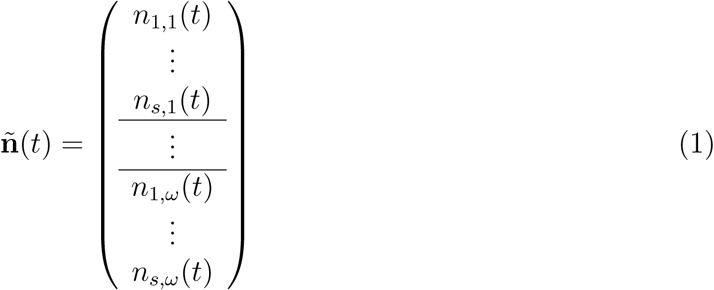

We use a projection interval of one day. In our data, no individual reproduced after 16 days, so we set both the maximum age (*ω*) and the maximum maternal age (*s*) to 16.

An individual with maternal age *i* and age *j* produces *f*_*ij*_ daughters in a day, and survives to age *j* + 1 with probability *p*_*ij*_. These vital rates are incorporated into a fertility matrix 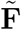 and a survival matrix **Ũ**. The population projection matrix **Ã**, which projects the population vector from one day to the next, is the sum of 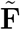 and **Ũ**, and the population dynamics are given by

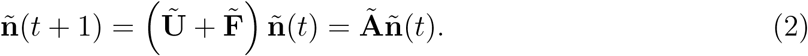

Caswell et al. [37] describe in detail the construction of **Ũ** and 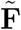 for general stage-by-age matrix models. In order to duplicate the results of our analyses, the reader needs:

**The matrix Ũ**. This block matrix takes the form

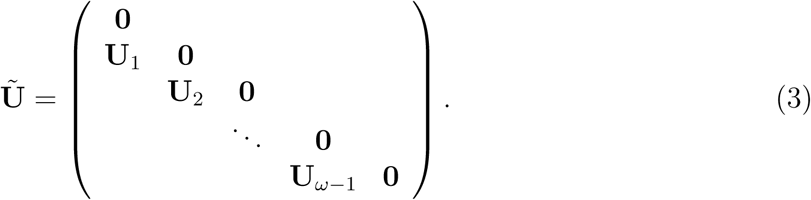

The **U**_*j*_ are *s* × *s* matrices with survival probabilities on the diagonal and zeros elsewhere:

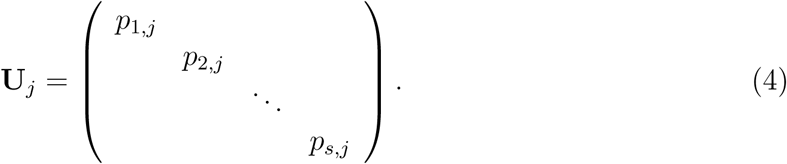

**The matrix** 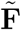. This matrix has a block first row composed of the *s* × *s* blocks **F**_*j*_:

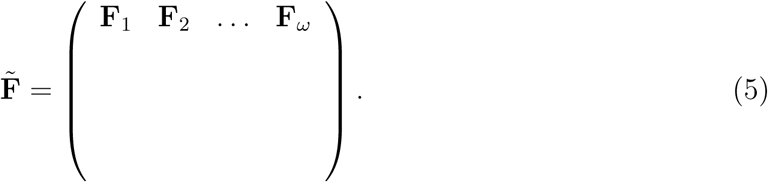

The block **F**_*j*_ is a fertility matrix for all females in age class *j*. Because the offspring of a mother of age *j* have maternal age *j*, the matrix ***F***_*j*_ contains zeros everywhere except in the *j*-th row, where the vector **f**_*j*_ = (*f*_1,*j*_, *f*_2,*j*_, …, *f*_*s,j*_) appears.

A population described by model (2) will converge to a stable structure 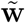 and grow exponentially at the rate log *λ*, where *λ* is the largest eigenvalue of the projection matrix **Ã** and 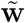 is the corresponding right eigenvector. We treat the intrinsic growth rate *λ* associated with a phenotype as a measure of the fitness of that phenotype [e.g. 2, 38, 39]. The selection gradient on a trait that affects 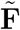or **Ũ** (e.g., *p*_*ij*_ or *f*_*ij*_) is the partial derivative of *λ* with respect to that trait [39]. Formulae for these selection gradients are given in [25].

## 3 Model parameterization

The fertilities *f*_*ij*_ and survival probabilities *p*_*ij*_ are the building blocks of model (2). We parameterize the dependence of these quantities on age and maternal age as follows.

To represent the dependence of fertility on age and maternal age, we adapted a model commonly used in human demography due to Coale and Trussell [40]. The model assumes that fertility is the product of a “natural fertility” that depends only upon age, and an additional factor that, beyond some fixed age, reduces the natural fertility. When fit to our laboratory data using maternal age as the additional factor (as described in the Supporting Information), the Coale-Trussell model produces a fertility schedule that increases sharply from two to four days of age. The schedules for individuals with different maternal ages diverge after age four. The fertility of individuals with older maternal age decline earlier and faster than the fertility of individuals with younger maternal age (Fig. 1A).

**Figure 1:**
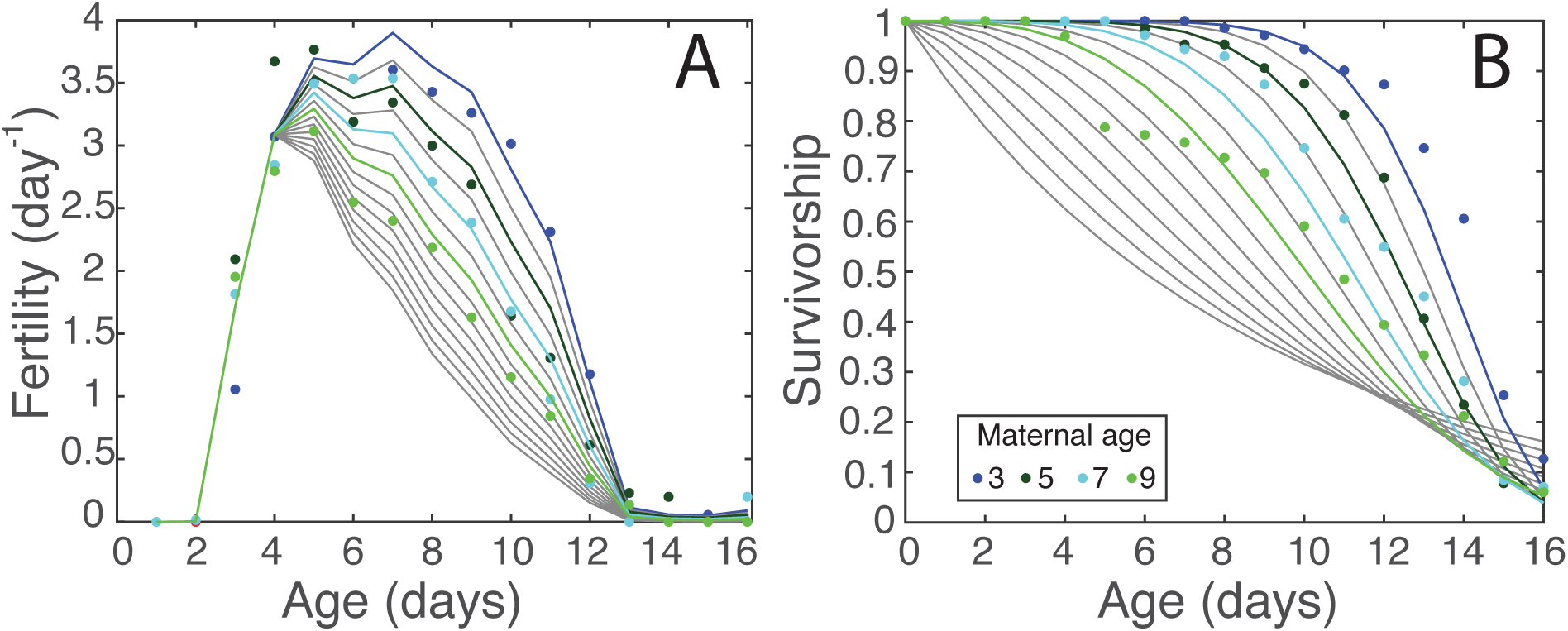
Fertility and survival schedules that underlie the projection matrix **Ã**. Observations are shown as filled circles, colored by their maternal age classification. Model fits to the data are shown as colored lines, and interpolated/extrapolated curves are shown in grey.

To represent the dependence of survival on age and maternal age we assumed that the distribution of the time to death within each maternal age is adequately described by the Weibull model [e.g., 41], with parameters that depend log-linearly on maternal age (see Supporting Information). The resulting best fit model predicts that survivorship decreases with maternal age for young individuals, but increases with maternal age for the very oldest individuals (Fig. 1B). These oldest individuals have extremely low fertility and are very rare in the population, so this old-age increase in survival with maternal age has negligible effect on our results.

### 3.1 Model modifications

The projection matrix **Ã** is estimated under laboratory culture conditions that lead to unrealistically high population growth rates for natural populations. This affects not only the growth rate, but also the population structure and selection gradients. To evaluate these effects, we created a hypothetical matrix 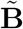 with reduced fertility, such as might result from resource limitation in nature, and repeated the calculations for this matrix. The fertilities in 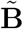 are obtained by dividing the fertilities in **Ã** by the net reproductive rate *R*_0_; this yields a stationary population with an intrinsic growth rate of 1. The survival schedule and the shape and timing of reproduction are unchanged.

The projection matrices **Ã** and 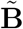 both include maternal effect senescence. To quantify the fitness costs of maternal effect senescence and to investigate the selective processes that could lead to its evolution, we need to compare **Ã** and 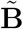 with hypothetical matrices in which offspring quality does not decline with increasing maternal age. We call these matrices **Ã** ^(*r*)^ and 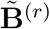, to indicate that maternal effect senescence has been removed. In **Ã** ^(*r*)^ and 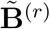, all individuals, regardless of maternal age, have the fertility and survival schedules corresponding to maternal age 3 in **Ã** and 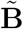, respectively. This maternal age group has the highest fertility rates and the largest survival probabilities. We then compute the selection gradients on **Ã** ^(*r*)^ and 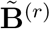, and also use them as reference matrices for performing an LTRE analysis.

## 4 Demographic and evolutionary analyses

The population projection matrices provide the link between the individual-level data from our laboratory experiments [16] and their ecological and evolutionary consequences for populations.

### 4.1 Fitness and population structure

The fitnesses of the four life histories described by each of the matrices are:

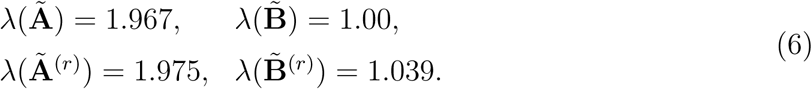

In the high-growth, maternal-age-senescent population, represented by **Ã**, the expected lifetime reproductive output (*R*_0_) is 22.42. The stable population structure is dominated by young individuals that were born to young mothers; individuals of age 1–3 d and with a maternal age of 1–5 d constitute 77% of the population (Fig. 2A). In the low-growth, maternal-age-senescent population, represented by 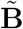, we have adjusted the fertility so that *λ* = 1 and *R*_0_ = 1. As a result, the stable population structure is much flatter; individuals of age 1–3 d and maternal age 1–5 d constitute only 11% of this population (Fig. 2B).

**Figure 2:**
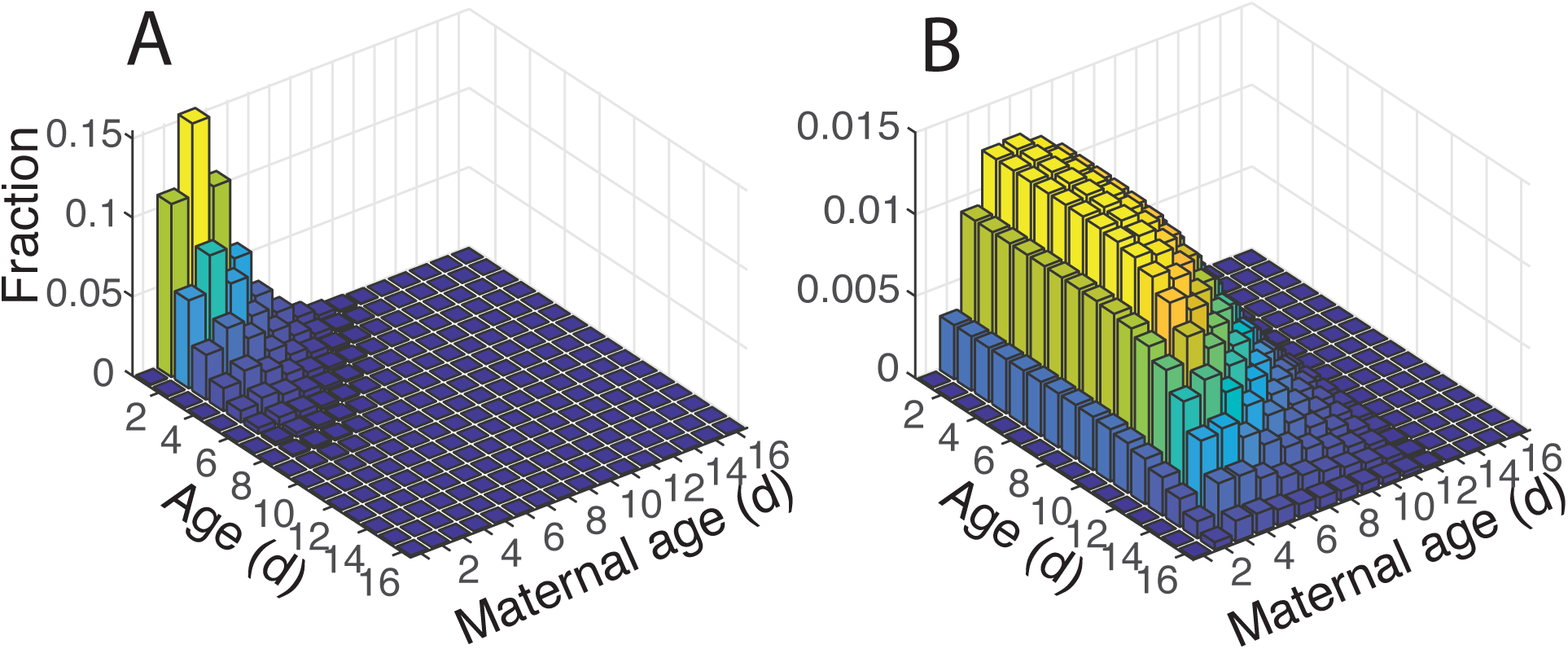
Stable age-by-maternal-age distributions for a high-growth population (**Ã**, panel A) and a low-growth population (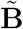, panel B), both with maternal age effect senescence. The height of each bar represents the percent of the stable population that is of that age and maternal age. The bars are colored by their height, corresponding with the values on the z-axis. Note the change in z-scale between the two panels.

Removing maternal effect senescence increases *λ* (equation (6)). The stable population structures are qualitatively unchanged (Fig. S3). The fitness differences due to maternal age senescence in high-growth laboratory conditions, as measured by the difference *λ*(**Ã**) − *λ*(**Ã** ^(*r*)^), is small (Δ*λ* = −0.008). Under those conditions almost all the population is at young ages and young maternal ages, where the effect of maternal age senescence is minimal. In the stationary population, the fitness difference 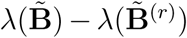 is larger than in the high-growth conditions (Δ*λ* = −0.039).

**Figure 3:**
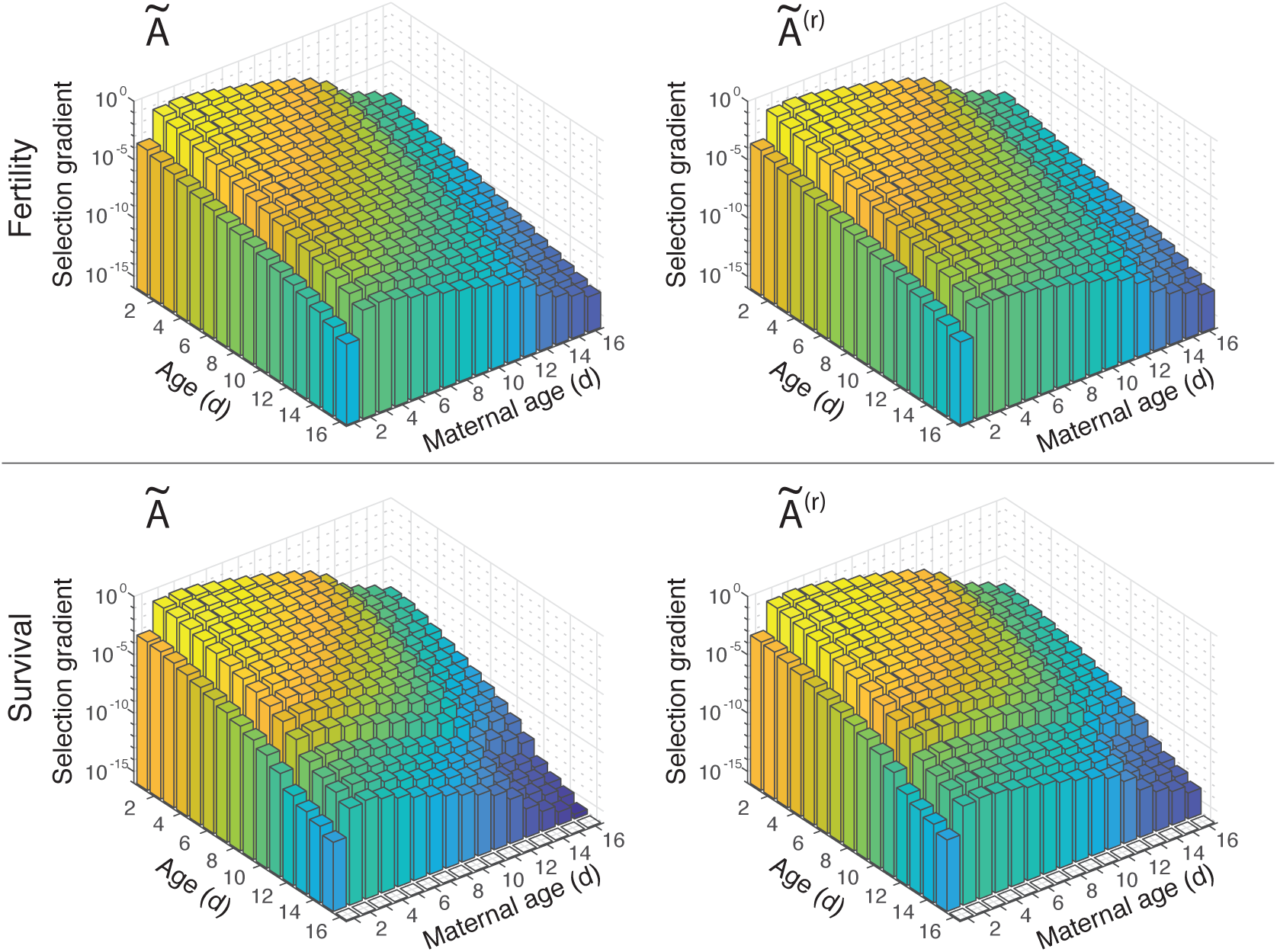
Selection gradients on survival and fertility in high-growth life histories, with and without the presence of maternal age effects. The panels in the top row are the selection gradients on fertility parameters, and the bottom row are for survival parameters. The left column shows the selection gradients in the high-growth environment with maternal effects, and the right column shows the selection gradients in the high-growth environment without maternal effects. The height of the bars is on a log-scale, and the bars are colored by their height, corresponding to the z-axis. All subpanels have the same z-axis limits and corresponding color axis.

### 4.2 Selection gradients and the evolution of maternal effect senescence

The strength of selection is measured by the selection gradients, defined as the derivatives of lambda with respect to survival or fertility [25]. The opportunity for selection to lead to senescence varies directly with the rate at which the selection gradient declines with age [42].

As expected [35], the selection gradients do not change qualitatively when growth rate is reduced, so we focus here on the (arguably more realistic) low-growth life histories; the high-growth results are found in Supplementary Information (Fig. 3).

The selection gradients on survival and fertility decline with age within any maternal age class (Fig. 4) as indeed they must, since individuals cannot change their maternal age group. As usual, the differences are large: 12 orders of magnitude or more (up to 15 orders of magnitude in the high-growth life histories). This decline provides the impetus for the evolution of age-related senescence.

**Figure 4:**
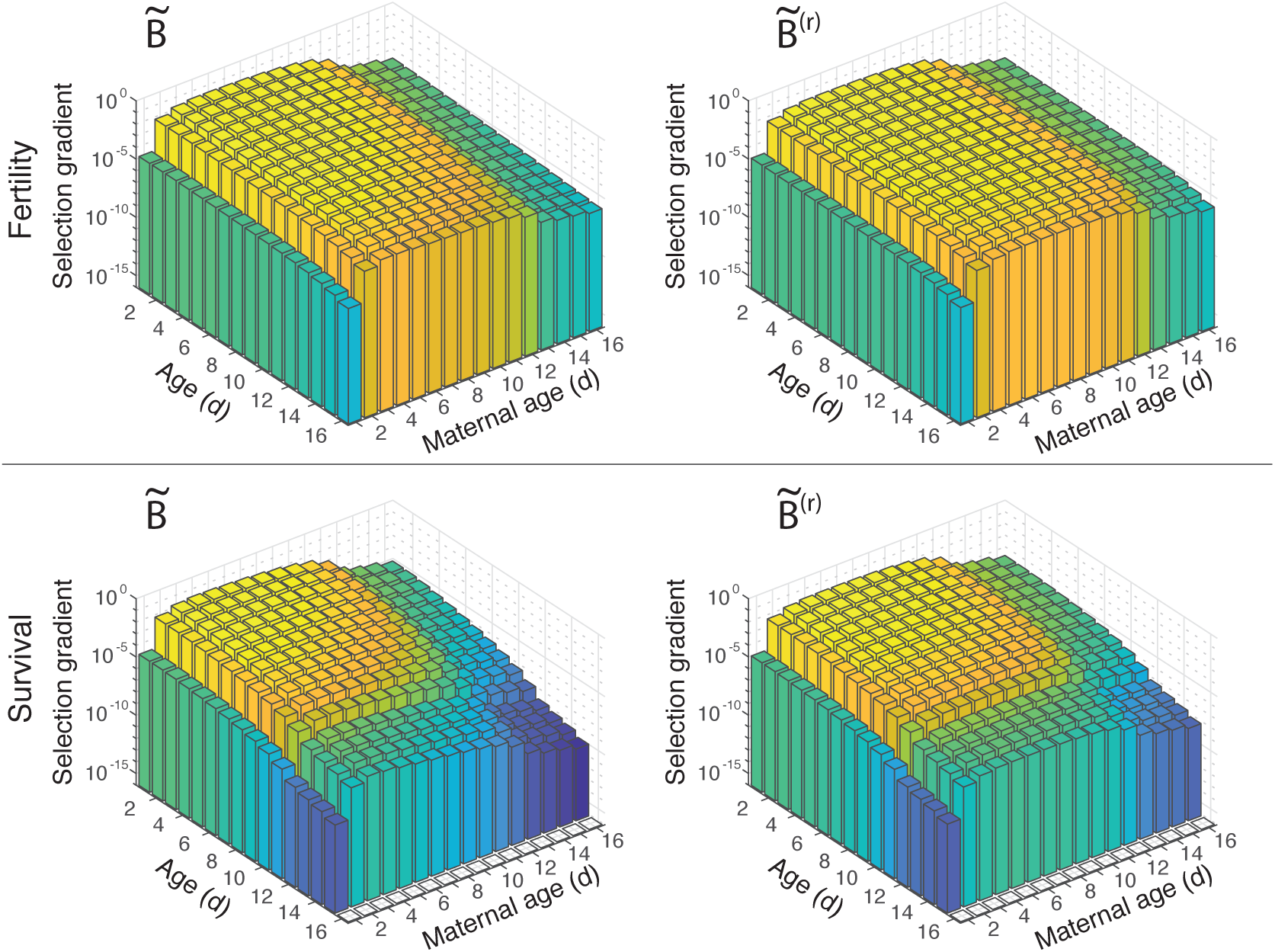
Selection gradients on survival and fertility in low-growth life histories, with and without the presence of maternal age effects. The panels in the top row are the selection gradients on fertility parameters, and the bottom row are for survival parameters. The left column shows the selection gradients in the low-growth environment with maternal effects, and the right column shows the selection gradients in the low-growth environment without maternal effects. The height of the bars is on a log-scale, and the bars are colored by their height, corresponding to the z-axis. All subpanels have the same z-axis limits and corresponding color axis.

Of central importance to our analysis, the selection gradients also decline with maternal age, after maternal age 3 for high-growth life histories, and after maternal age 4 or 5 for low-growth life histories. (Fig. 4). The declines in the selection gradients after their initial increase are large: 5–7 orders of magnitude in high-growth conditions and 2-3 orders of magnitude in low-growth conditions. That is, a unit reduction in survival or fertility at old maternal ages can be paid for by a much smaller increase, one hundreth to one millionth as large, at younger maternal ages.

This result implies that maternal effect senescence can evolve in the same way as age senescence. Traits that reduce survival or fertility of the offspring of older mothers can be balanced by much smaller improvements at younger maternal ages. Even in the absence of such pleiotropic effects, selection will be less efficient at removing deleterious mutations at older ages, permitting them to accumulate.

The matrices **Ã** and 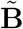 already contain maternal effect senescence, and so the selection gradients calculated from them apply, strictly speaking, to increases in already existing maternal effects. In the life histories described by **Ã** ^(*r*)^ and 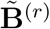, maternal age effects have been removed. The selection gradients, on both survival and fertility, still decline with increasing maternal age, implying that selection not only favors maternal effect senescence when it is already present in *B. manjavacas*, but that it can also easily arise de novo.

### 4.3 The sources of fitness differences: LTRE analysis

The fitness differences due to maternal effect senescence are measured by the difference in *λ* between **Ã** and **Ã** ^(*r*)^, or between 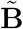 and 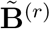, in high-growth and low-growth environments, respectively. The differences in *λ* are consequences of all the differences in all the vital rates at all combinations of age and maternal age. The contributions of each of these differences to the difference in fitness is calculated using an LTRE analysis [35, Chap. 10].

An LTRE analysis decomposes the difference between the population growth rates of two projection matrices into the contributions from the differences in the entries in those matrices. To focus on the contributions from the maternal age groups, and to separate those contributions into survival and fertility, we summed all the contributions over age within each maternal age group.

In the high-growth population, the fitness difference due to maternal effect senescence is Δ*λ* = −0.008. The LTRE analysis reveals that this change in fitness arises mostly from decreases in fertility; the contributions of these decreases are much larger than the contributions from the decreases in survival (Fig. 5A). The greatest contribution to the fitness difference comes from the decrease in fertility of individuals with maternal age between 4 and 6 d. The fitness differences due to changes in survival associated with maternal effect senescence are small, and peak at maternal ages 4–6 d.

**Figure 5:**
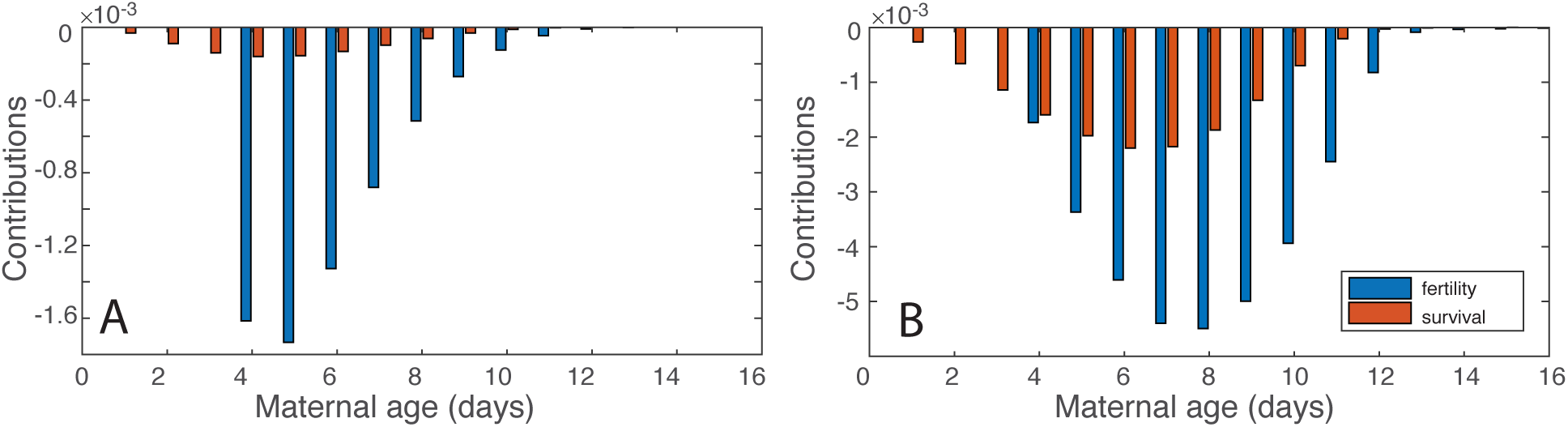
Results of the Life Table Response Experiments in the high-growth population (comparing **Ã** with **Ã**^(*r*)^, panel A) and the low-growth population (comparing 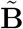 with 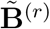, panel B). The contributions of fertility and survival, summed across ages, show the effect on *λ* from changes in fertility and survival for offspring from a given maternal age. Note the change in scale between the panels.

In the low-growth population, the fitness difference is Δ*λ* = −0.039, which is 5 times larger in magnitude than the difference in the high-growth population. As in the high-growth population, the fitness difference arises mostly from decreased fertility, but decreased survival makes a larger contribution than in the high-growth population. The largest contributions at maternal ages 7–9 d. The contributions from survival peak at maternal age 6–7 d (Fig. 5B).

Taken together, our LTRE analyses show that the decreased fitness resulting from maternal effect senescence is primarily caused by decreased lifetime fertility of offspring, and primarily at early maternal ages (3-5 d) for the high-growth population with a shift towards middle life (6-9 d) for the low-growth population.

## 5 Discussion

Maternal effect senescence, defined as a decrease in offspring survival and/or reproduction with increasing maternal age, has been reported in our study organism, the rotifer *B. manjavacas* [16, 22]. In this study, we quantified the population and evolutionary consequences of these maternal age effects, by incorporating them into a multistate matrix population model. We found that age-specific survival and fertility decrease with increasing maternal age, and that this carries a cost in fitness compared to a hypothetical life cycle in which the maternal effect senescence has been removed. The cost is obscured in the luxurious (for a rotifer) laboratory conditions, because the unrealistic population growth rates lead to a population structure in which almost all individuals are young offspring of young mothers. The cost is larger in a life cycle in which the population is rendered stationary by reducing fertility.

The decline in selection gradients provides the opportunity for the evolution of maternal effect senescence. LTRE analysis demonstrates that the population-level fitness differences are primarily due to decreased fertility of individuals with early to intermediate maternal ages.

In a previous theoretical study, Moorad and Nussey [7] found that reproductive senescence and maternal effect senescence can evolve separately. They take a quantitative genetics approach, and maternal effect senescence only appears as a decrease in neonatal survival with increasing maternal age. Our study complements theirs by using experimental data to estimate the parameters in a demographic model. Furthermore, maternal effect senescence in our study reduces survival and fertility throughout the life of the offspring of older mothers.

Consistent with Hamilton’s theory of senescence, we find that selection gradients decrease with increasing maternal age, suggesting the two mechanisms by which senescence can evolve—antagonistic pleiotropy and mutation selection balance—could also lead to the evolution of maternal effect senescence.

Importantly, the pattern of decreasing selection gradients with increasing maternal age is seen in both the high-growth (**Ã**) and low-growth 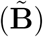 populations, demonstrating that our findings are relevant to natural populations, and are not artifacts of the luxurious laboratory environment. The opportunity for the evolution of maternal effect senescence persists even when we decrease fertility to simulate resource limitation. Furthermore, when we remove maternal effect senescence from both the high-growth (**Ã**) and low-growth populations 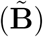, the pattern of decreasing selection gradients persists, suggesting that maternal effect senescence may evolve de novo.

The potential ease with which maternal effect senescence can arise raises questions about why it is not observed in all species. For example, the extremely diverse group of teleost fishes generally have increased fitness with advanced maternal age; not only do older mothers produce more eggs, but those eggs have a higher likelihood of hatching and of surviving to maturity [43]. In contrast to rotifers, fish have indeterminate growth and an increasing fertility schedule. Larger females make a greater investment in offspring, so demographic models of fish are generally size-structured as opposed to age-structured. Conversely, rotifers demonstrate determinate growth, with cell number fixed at the time of hatching.

Stage-classified models can lead to non-monotonic selection gradient patterns that differ significantly from those presented in this paper. For example, plant life cycles can vary significantly from animal life cycles, with individuals moving through life stages or size classes in more complex patterns. In an analysis of age-by-size structured models of plants, selection gradients did not uniformly favor the evolution of senescence, but in fact at some ages operated against the evolution of senescence [25]. Within stages (size, sex, and maturity), selection gradients increased with age until an intermediate “critical age” before declining again. This demonstrates that when other types of traits are accounted for in a multi-state model, the expected patterns of age senescence may not hold.

In organisms with life histories different from that of *B. manjavacas*, we may see distinct patterns of maternal effect senescence. For example, organisms that exhibit batch rather than continuous reproduction or a different degree of maternal investment in reproduction might be expected to show different selection gradients. Maternal effect senescence may also evolve differently in taxa with life cycles with complex stage transitions such as metamorphosis or sharp changes in survivorship due to habitat transitions. Additional types of maternal effects, such as imprinting or transgenerational epigenetic inheritance due to maternal experiences with specific environmental conditions (e.g., the Dutch Hunger Winter and Quebec Ice Storm [44, 45]) could complicate our understanding of maternal effect senescence.

Individual vital rates are determined by a myriad of life history parameters or individual states. In this study, we explored an age-by-stage model where the stage–maternal age– of each individual was fixed at birth. However, stage can also represent non-fixed traits such as size, sex, maturity state, or combinations of these [25]. The techniques for building and analyzing matrix models with two individual states are well-developed, and the stage classifications used are flexible. The mathematical equations to operate on three or more state dimensions are complicated [46], but may enable the exploration of more complex trait combinations, such as exploring the fitness effects of maternal age across multiple generations or in varied environments.

In this study we quantified the demographic and fitness consequences of maternal effect senescence, and the evolutionary conditions that may allow it to arise. For our model population of rotifers, in both high-growth and low-growth environments, and with maternal effects both present and absent, we find patterns of selection that conform to Hamilton’s theory for the evolution of senescence, namely that selection gradients decrease at advanced ages and maternal ages. This is important, as it demonstrates that maternal effect senescence can evolve even though the populations exhibiting maternal effect senescence experience decreased fitness. Future work will determine how maternal effect senescence impacts individual-level variance in expected lifespan and reproductive output, and how maternal effect senescence determines fitness across multiple generations. Valuable insights can gained by using the flexible model framework presented here to analyze maternal effect senescence in taxa with different reproductive strategies and fertility schedules.

## 6 Methods

### Notation

Matrices are denoted by bold uppercase symbols (e.g., **U**), vectors by bold lowercase symbols (e.g., **w**). Symbols with a tilde (e.g.,**ñ**) are block-structured, with maternal age classes grouped within ages.

### Lifetable Experiments

We conducted lifetable experiments using the Russian strain of the monogonont rotifer *Brachionus manjavacas* (BmanRUS) as previously described [16]. To avoid residual undefined parental effects in experimental populations, we synchronized the maternal ages of the great-grand-maternal and grand-maternal generations for the experimental maternal (F0) cohort by collecting eggs from 3 – 5 d old females for two generations. To obtain the F0, eggs from the age-synchronized grand-maternal culture were harvested by vortexing and allowed to hatch over 16 hours. Neonates were deposited individually into 1 ml of 15 ppt seawater and 6*x*10^5^ cells/ml of the chlorophyte algae, *Tetraselmis suecica*, in wells of 24-well tissue culture plates (n=187). To obtain the F1 cohorts F1_3_, F1_5_, F1_7_, F1_9_, at maternal ages of 3, 5, 7, and 9 d, respectively, we isolated one female neonate hatched within the previous 24 h per F0 female. F1 neonates were placed individually in wells of 24-well plates with 1 ml seawater and algae as above (n = 72 for each F1 cohort). Every 24 h, we recorded survival, reproductive status (whether carrying eggs), and the number of live offspring for each F0 and F1 individual; the female was then transferred to a new well with fresh algae of known concentration. Survivorship data were right censored in cases where individuals were lost prior to death.

## Acknowledgements

This work was supported by grant 5K01AG049049 from the National Institute on Aging to KEG, and by the Bay and Paul Foundations. HC and SvD were supported by the European Research Council through Advanced Grants 322829 and 788195, and by the Dutch Research Council through grant ALWOP.2015.100. CMH was supported by an NSF Graduate Research Fellowship, and the J. Seward Johnson Endowment in support of the Woods Hole Oceanographic Institution’s Marine Policy Center.

## 7 Supporting Information

### 7.1 Fertility model

The Coale and Trussell model [40] in human demography represents differences in the fertility schedules of populations by using a ‘natural fertility schedule’ 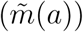, an age at which the population-specific schedules can diverge from the natural fertility schedule (*a*_0_), and a parameter that controls how rapidly the schedules diverge (*θ*). The model is

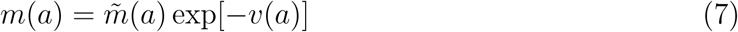

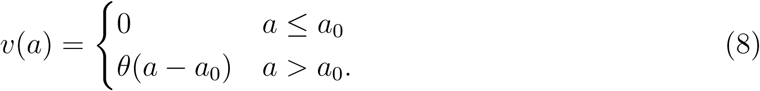

The Coale and Trussell model is typically fit to each population separately by choosing *θ* and *a*_0_ to satisfy some statistical criterion (e.g., minimizing the sum of squared deviations between the model and the data).

If we had observed the fertility schedules of individuals in every maternal age class, we could have used model (7)–(8) directly. Our model, however, needs the fertility schedule for maternal ages that we did not measure in the laboratory experiment. We therefore amended the Coale and Trussell model to incorporate the effects of maternal age *g*:

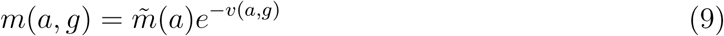

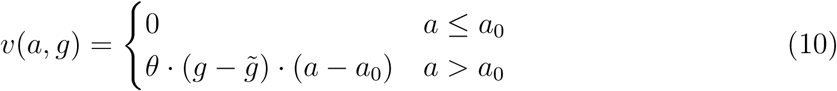

We now imagine that the “natural fertility schedule” is associated with a “natural maternal age” which, in model (9)–(10), is the new parameter 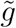.

To fit model (9)–(10), we minimized the sum of squared deviations between the model and the data over all of the possible choices of

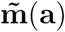. We assumed that natural fertility schedule 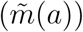 was one of five functions: one of the four observed fertility schedules for mothers with maternal age equal to 3, 5, 7 or 9; or the average fertility schedule over all maternal ages.

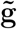. With each of the five candidate natural fertility schedules, we associate exactly one value of 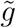. For the four observed schedules, with maternal ages 3, 5, 7, and 9, we set 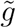 equal to the maternal age. For the fifth, average, candidate schedule we set 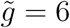.

**a**_**0**_. Based on visual inspection of the observed fertility schedules (Fig. 1), we assumed that *a*_0_ ∈ {2, 3, 4, 5}.

*θ*. The rate parameter *θ* was assumed to be a real positive number.

The best fit is acheived by the ‘average’ fertility schedule for 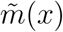, with 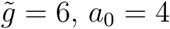, and *θ* = 0.0192. The data, model fits, and extrapolated curves are shown in the main text (Fig. 1A).

### 7.2 Time to death Weibull

The Weibull model is a common representation of the time-to-failure in survival probability estimation for living organisms. It is a continuous probability distribution described by two parameters, and the shape of the probability density function (*pdf*) is very flexible. From the individual culture experiments described in [47], we have the time-to-death of many individual rotifers—with maternal ages 3, 5, 7, and 9 days—and we used this data to fit the Weibull distribution:

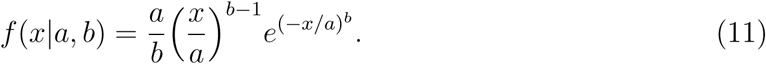

The failure time, *x* of each rotifer in the dataset is the oldest age (days) at which it was observed alive. The Weibull *pdf* (Equation 11) is fit to observed failure times in each maternal age cohort, yielding 4 Weibull models. Inspection of the parameters from those 4 models indicates that the parameters *a* and *b* vary log-linearly with maternal age group (Fig. 6), such that the Weibull parameters are themselves parametric functions of maternal age group, *g*. Although a linear model would fit equally well to the 4 points (Figs. 6), the log-linear form is necessary so that the Weibull parameters will both be non-negative for maternal age groups up to *g* = 16. With this formulation, the time-to-death description of mortality probabilities is a function of a single variable, *g*, which varies among groups, and 4 parameters that are defined for the population as a whole (the slope and intercept of the log-linear functions for *a* and *b*). The resulting Weibull *pdf* s are shifted towards earlier and more evenly spread times of death as *g* increases (Fig. 7). The resulting survivorship curves are shown in the main text (Fig. 1B).

**Figure 6:**
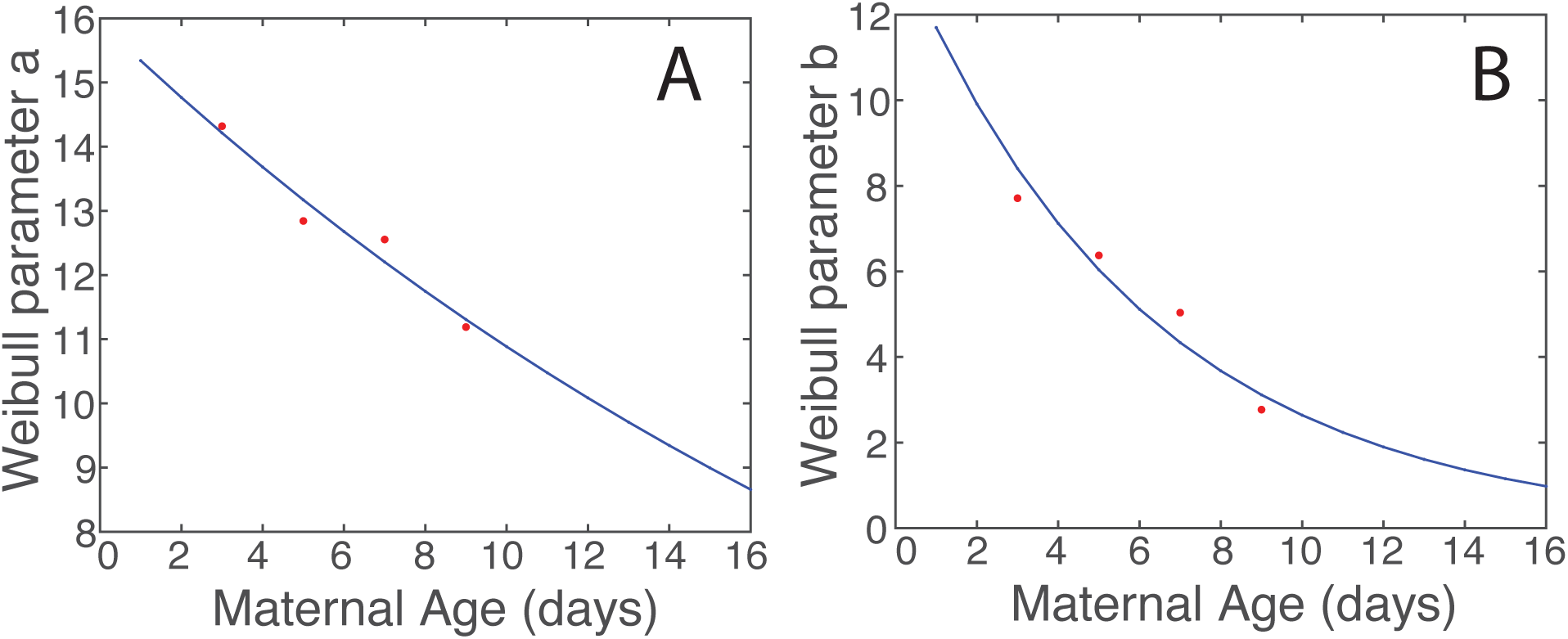
Weibull model parameters *a* (panel A) and *b* (panel B) vary log-linearly with maternal age group. Red points represent the observed Weibull parameters for maternal ages 3, 5, 7, and 9. The blue lines are the log-linear best fits, *a* = *exp*(−0.038 · *g* + 2.77) and *b* = *exp*(−0.165 *g* + 2.63).

**Figure 7:**
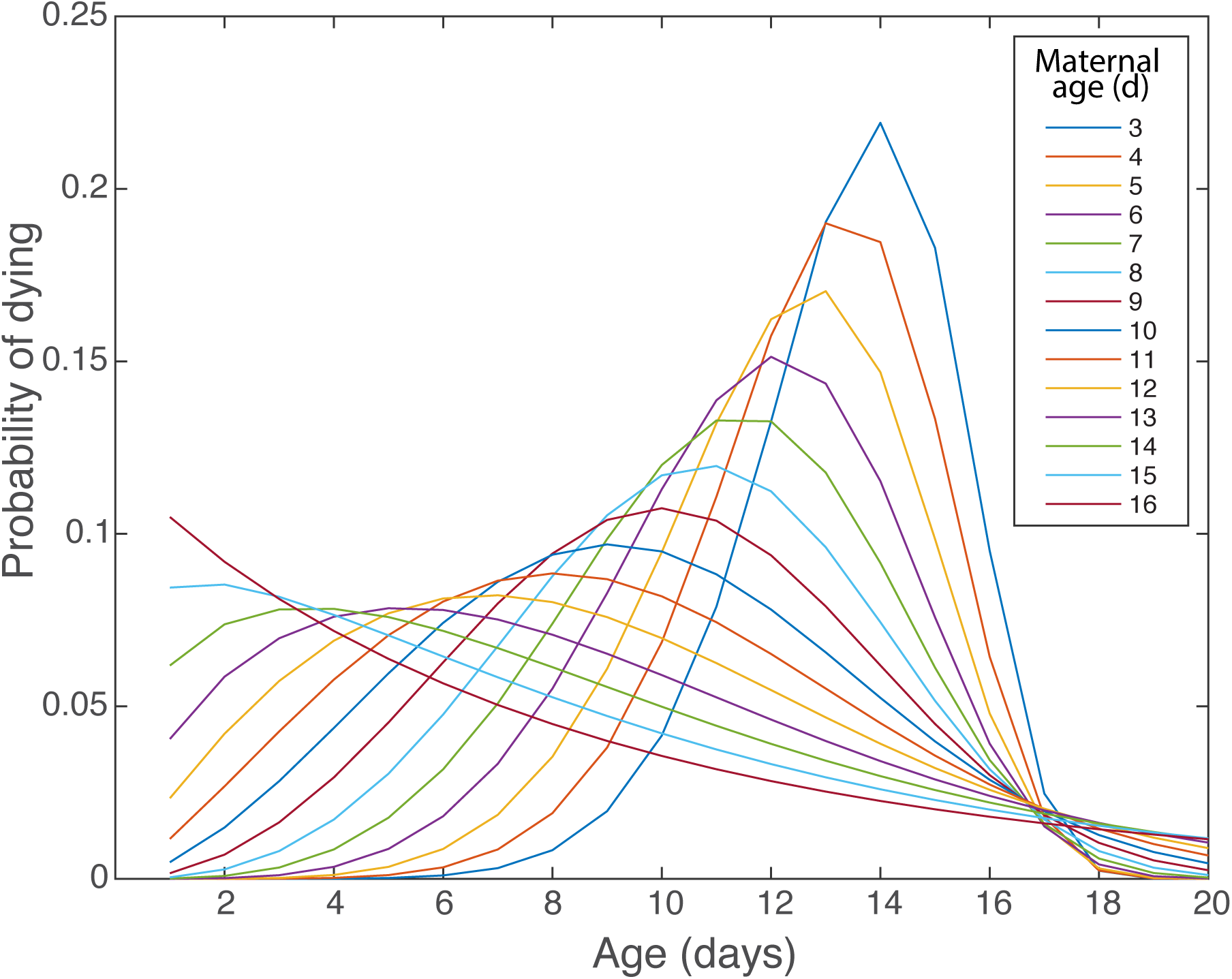
Weibull probability density functions from the full model that varies with *g*, for *g* ∈ [3, 16]. The blue curve with the highest peak is for maternal age group 3, and the red curve that is highest at age 1 day is for maternal age group 16.

**Figure 8:**
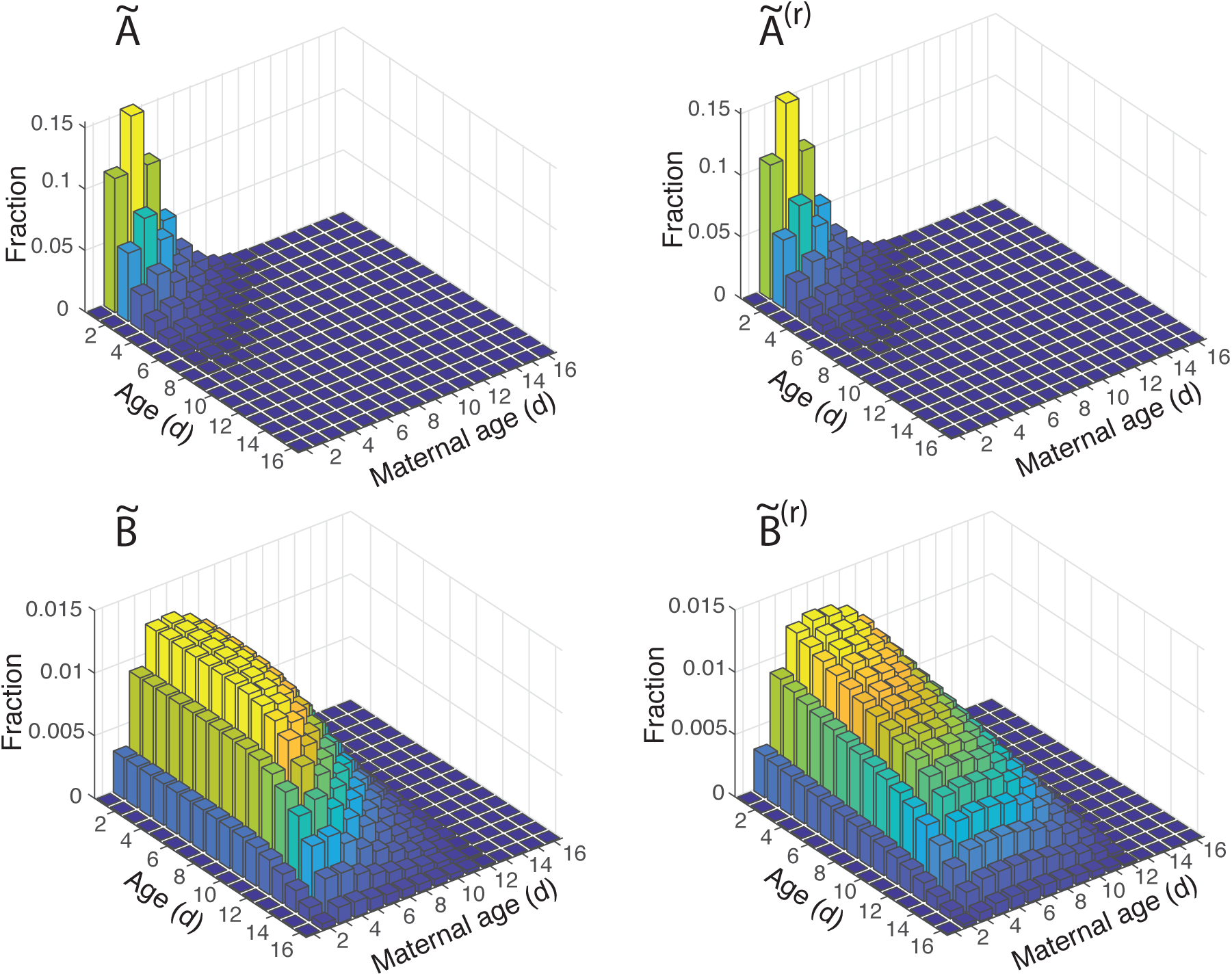
Stable age-by-maternal-age distributions for all four population models: high-growth population with maternal age effects (**Ã**, top left), high-growth population without maternal age effects (**Ã** ^(*r*)^, top right), low-growth population (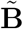, bottom left) with maternal age effects, and low-growth population without maternal age effects (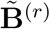, bottom right). The height of each bar represents the percent of the stable population that is of that age and maternal age. The bars are colored by their height, corresponding with the values on the z-axis. The upper two panels (**Ã** and **Ã** ^(*r*)^) have the same z-axis and colorbar, which differs from the z-axis and colorbar for the bottom two panels (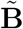 and 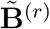).

### 7.3 Birthflow model

Birth and death for the lab-cultured rotifers are continuous processes, and by censusing once per day, information on the age of an individual is limited. Neonates were assigned an age of 0 on the day that they were first counted in the census, but their true ages were between 0 and 1. Therefore, estimates of both fecundity and survival probability represent averages over the census interval (1 day), and a birth-flow model must be used to build up the transition matrix, **Ã** [35].

### 7.4 Survivorship curves

The survivorship curve (the proportion of neonates that survive to each age) was calculated as the complementary cumulative distribution function for the Weibull model. The transition probabilities for the matrix model formulation are given in Equation 12, where *P*_*i*_ is the probability of surviving from age *i* to age *i* + 1, and *l* is the survivorship curve derived from the Weibull model.

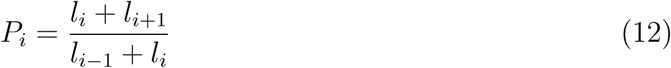

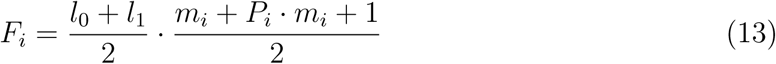

Likewise, the observed number of offspring produced by an individual in an interval averages over several processes that are unobserved during the interval. Production of neonates by a given age class, *F*_*i*_, is determined by the fertility at that age, the fertility at the next age, the probability that individuals survive to that next age, and the probability that the offspring survive long enough to be counted (Equation 13). The average offspring must survive 0.5 days to be counted, and the fertility of the mother is an average, weighted by the probability of surviving to the next age [35]. The *m* values in Equation 13 come directly from the fertility model described in Section 7.1.

## References

[1] P.B. Medawar. An unsolved problem in biology. H. K. Lewis, London, England, 1952.

[2] William D Hamilton. The moulding of senescence by natural selection. Journal of Theoretical Biology, 12(1):12–45, 1966.

[3] Daniel Fabian and Thomas Flatt. The evolution of aging. Nature Education Knowledge, 3(10):9, 2011.

[4] Michael R Rose. Evolutionary Biology of Aging. Oxford University Press, Oxford, U.K., 1990.

[5] Owen R Jones, Alexander Scheuerlein, Roberto Salguero-Gómez, Carlo Giovanni Camarda, Ralf Schaible, Brenda B Casper, Johan P Dahlgren, Johan Ehrlén, María B García, Eric S Menges, Pedro F Quintana-Ascencio, Hal Caswell, Annette Baudisch, and James W Vaupel. Diversity of ageing across the tree of life. Nature, 505(7482): 169–173, Jan 2014. doi: 10.1038/nature12789.

[6] Richard P Shefferson, Roberto Salguero-Gómez, and Owen R Jones editors. The Evolution of Senescence in the Tree of Life. Cambridge University Press, 2017.

[7] J. A. Moorad and D. H. Nussey. Evolution of maternal effect senescence. Proc Natl Acad Sci U S A, 113(2):362–7, 2016. ISSN 1091-6490 (Electronic) 0027-8424 (Linking). doi: 10.1073/pnas.1520494113. URL https://www.ncbi.nlm.nih.gov/pubmed/26715745.

[8] Albert I Lansing. A transmissible, cumulative and reversible factor in aging. J Gerontol, 2(3):228–239, 1947.

[9] R. de la Fuente-Fernandez. Maternal effect on parkinson’s disease. Ann Neurol, 48(5): 782–7, 2000. ISSN 0364-5134 (Print) 0364-5134.

[10] M.J. Hercus and A.A. Hoffman. Maternal and grandmaternal age influence offspring fitness in drosophila. Proc R Soc Lond B Biol Sci, 267:2105–2110, 2000. doi: 10.1098/rspb.2000.1256.

[11] W. Fox, Charles M.L. Bush, and W.G. Wallin. Maternal age affects offspring lifespan of the seed beetle, callosobruchus maculatus. Funct Ecol, 17:811–820, 2003. doi: 10.1111/j.1365-2435.2003.00799.x.

[12] Nicole M. Talge, Charles Neal, Vivette Glover, Translational Research the Early Stress, and Prevention Science Network. Antenatal maternal stress and long-term effects on child neurodevelopment: how and why? J Child Psychol Psychiatry, 48:246–261, 2007. doi: 10.1111/j.1469-7610.2006.01714.x.

[13] S. Bouwhuis, O. Vedder, and P. H. Becker. Sex-specific pathways of parental age effects on offspring lifetime reproductive success in a long-lived seabird. Evolution, 69(7):1760–71, 2015. ISSN 1558-5646 (Electronic) 0014-3820 (Linking). doi: 10.1111/evo.12692. https://www.ncbi.nlm.nih.gov/pubmed/26095174.

[14] J. Schroeder, S. Nakagawa, M. Rees, M. E. Mannarelli, and T. Burke. Reduced fitness in progeny from old parents in a natural population. Proc Natl Acad Sci U S A, 112(13):4021–5, 2015. ISSN 1091-6490 (Electronic) 0027-8424 (Linking). doi: 10.1073/pnas.1422715112. https://www.ncbi.nlm.nih.gov/pubmed/25775600.

[15] M. C. Bloch Qazi, P. B. Miller, P. M. Poeschel, M. H. Phan, J. L. Thayer, and C. L. Medrano. Transgenerational effects of maternal and grandmaternal age on offspring viability and performance in drosophila melanogaster. J Insect Physiol, 100:43–52, 2017. ISSN 1879-1611 (Electronic) 0022-1910 (Linking). doi: 10.1016/j.jinsphys.2017.05.007. https://www.ncbi.nlm.nih.gov/pubmed/28529156.

[16] Martha J. Bock, George C. Jarvis, Emily L. Corey, Emily E. Stone, and Kristin E. Gribble. Maternal age alters offspring lifespan, fitness, and lifespan extension under caloric restriction. Scientific Reports, 9(3138), 2019. doi: 10.1038/s41598-019-40011-z.

[17] T. G. Benton, J. J. St Clair, and S. J. Plaistow. Maternal effects mediated by maternal age: from life histories to population dynamics. J Anim Ecol, 77(5):1038–46, 2008. ISSN 1365-2656 (Electronic) 0021-8790 (Linking). doi: 10.1111/j.1365-2656.2008.01434.x. http://www.ncbi.nlm.nih.gov/pubmed/18631260.

[18] M. C. Bloch Qazi, P. B. Miller, P. M. Poeschel, M. H. Phan, J. L. Thayer, and C. L. Medrano. Transgenerational effects of maternal and grandmaternal age on offspring viability and performance in drosophila melanogaster. J Insect Physiol, 100:43–52, 2017. ISSN 1879-1611 (Electronic) 0022-1910 (Linking). doi: 10.1016/j.jinsphys.2017.05.007. https://www.ncbi.nlm.nih.gov/pubmed/28529156.

[19] Stefanie Kern, Martin Ackerman, Stephen C. Stearns, and Tadeusz J. Kawecki. Decline in offspring viability as a manifestation of aging in drosophila melanogaster. Evolution, 55(9):1822–1831, 2001.

[20] Alexander Graham Bell. The duration of life and conditions associated with longevity: Study of the Hyde geneology. Genealogical Record Office, Washington, D.C., 1918.

[21] Lindsay Farrer, Adrienne Cupples, Dan K. Kiely, P. Michael Conneally, and Richard H. Myers. Inverse relationship between age at onset of huntington disease and paternal age suggests involvement of genetic imprinting. American Journal of Human Genetics, 50: 528–535, 1992.

[22] Kristin E Gribble, George Jarvis, Martha Bock, and David B Mark Welch. Maternal caloric restriction partially rescues the deleterious effects of advanced maternal age on offspring. Aging Cell, 13(4):623–630, Aug 2014. doi: 10.1111/acel.12217.

[23] S. J. Plaistow, C. Shirley, H. Collin, S. J. Cornell, and E. D. Harney. Offspring provisioning explains clone-specific maternal age effects on life history and life span in the water flea, daphnia pulex. Am Nat, 186(3):376–89, 2015. ISSN 1537-5323 (Electronic) 0003-0147 (Linking). doi: 10.1086/682277. http://www.ncbi.nlm.nih.gov/pubmed/26655355.

[24] M. F. Perez and B. Lehner. Intergenerational and transgenerational epigenetic inheritance in animals. Nat Cell Biol, 21(2):143–151, 2019. ISSN 1476-4679 (Electronic) 1465-7392 (Linking). doi: 10.1038/s41556-018-0242-9. https://www.ncbi.nlm.nih.gov/pubmed/30602724.

[25] Hal Caswell and Roberto Salguero-Gómez. Age, stage and senescence in plants. J Ecol, 101(3):585–595, May 2013. doi: 10.1111/1365-2745.12088.

[26] John J. Gilbert. Rotifer ecology and embryological induction. Science, 151:1234–1237, 1966.

[27] John J. Gilbert. Asplancha and posterolateral spine induction in brachionus calyciflorus. Arch. Hydrobiol., 64:1–62, 1967.

[28] John J. Gilbert. Susceptibilities of ten rotifer species to interference from daphnia pulex. Ecology, 69:1826–1838, 1988.

[29] John J. Gilbert. Kairomone-induced morphological defenses in rotifers. In Ralph Tollrian and C. Drew Harvell, editors, The ecology and evolution of inducible defenses, pages 127–141. Princeton University Press, Princeton, NJ, 1999.

[30] John J. Gilbert. Specificity of crowding response that induces sexuality in the rotifer brachionus. Limnology and Oceanography, 48(3):1297–1303, 2003.

[31] John E. Havel and Stanely I. Dodson. Chaoborus predation on typical and spined morphs of daphnia pulex: Behavioral observations. Limnology and Oceanography, 29 (3):487–494, 1984.

[32] Donald A. Krueger and Stanely I. Dodson. Embryological induction and predation ecology in daphnia pulex. Limnology and Oceanography, 26(2):219–223, 1981.

[33] Terry W Snell, Rachel K Johnston, Kristin E Gribble, and David B Mark Welch. Rotifers as experimental tools for investigating aging. Invertebr Reprod Dev, 59(1):5–10, 2015. doi: 10.1080/07924259.2014.925516.

[34] Kristin E. Gribble and Terry W. Snell. Rotifers as a model for the biology of aging. In Jeffrey L Ram and P. Michael Conn, editors, Conn’s Handbook of Models for Human Aging, chapter 36, pages 483–495. Academic Press, London, U.K., 2nd edition, 2018. ISBN 978-0-12-811353-0.

[35] Hal Caswell. Matrix Population Models: Construction, Analysis, and Interpretation. Sinauer Associates, 2001.

[36] Hal Caswell. Sensitivity Analysis: Matrix Methods in Demography and Ecology. Springer, 2019.

[37] Hal Caswell, Charlotte de Vries, Nienke Hartemink, Gregory Roth, and Silke F van Daalen. Age × stage-classified demographic analysis: a comprehensive approach. Ecol Monogr, 88(4):560–584, Nov 2018. doi: 10.1002/ecm.1306.

[38] Johan A J Metz, Roger M Nisbet, and Stefan A H Geritz. How should we define ‘fitness’ for general ecological scenarios? Trends in Ecology and Evolution, 7(6):198–202, 1992.

[39] Russell Lande. A quantitative genetic theory of life history evolution. Ecology, 63(3): 607–615, 1982.

[40] Ansley J Coale and T James Trussell. Model fertility schedules: variations in the age structure of childbearing in human populations. Population Index, pages 185–258, 1974.

[41] D.R. Cox and D. Oakes. Analysis of Survival Data. Number 21 in Monographs on Statistics and Applied Probability. Chapman & Hall, London, UK, 1984.

[42] Hal Caswell and Esther Shyu. Senescence, selection gradients and mortality. In Richard P Shefferson, Owen R Jones, and Roberto Salguero-Gómez, editors, The Evolution of Senescence in the Tree of Life, pages 56–82. Cambridge University Press, 2017.

[43] Mark a Hixon, Darren W Johnson, and Susan M Sogard. BOFFFFs: on the importance of conserving old-growth age structure in fishery populations. ICES Journal of Marine Science, 71(8):2171–2185, oct 2014. ISSN 1054-3139. doi: 10.1093/icesjms/fst200. https://academic.oup.com/icesjms/article-lookup/doi/10.1093/icesjms/fst200.

[44] Tessa Roseboom, Susanne de Rooij, and Rebecca Painter. The dutch famine and its long-term consequences for adult health. Early Human Development, 82:485–491, 2006. doi: 10.1016/j.earlhumdev.2006.07.001.

[45] Lei Cao-Lei, Renaud Massart, Matthew J. Suderman, Ziv Machnes, Guillaume Elgbeili, David P. Laplante, Moshe Szyf, and Suzanne King. Dna methylation signatures triggered by prenatal maternal stress exposure to a natural disaster: Project ice storm. PLoS ONE, 9(9):e107653., 2014. doi: 10.1371/journal.pone.0107653.

[46] Gregory Roth and Hal Caswell. Hyperstate matrix models: extending demographic state spaces to higher dimensions. Methods in Ecology and Evolution, 7(12):1438–1450, 2016.

[47] Kristin E Gribble and David B Mark Welch. Genome-wide transcriptomics of aging in the rotifer brachionus manjavacas, an emerging model system. BMC Genomics, 18(1): 217, 03 2017. doi: 10.1186/s12864-017-3540-x.

